# REverse-transcriptase ACTivity with CRISPR (REACTR) Assay for Rapid and User-Friendly Therapeutic Drug Monitoring in Cytomegalovirus Care

**DOI:** 10.64898/2025.12.08.693028

**Authors:** Willow A. Chernoske, Maya A. Singh, Carrie H. Lin, Megan M. Chang, Cosette A. Craig, Albert H. Park, Joseph E. Rower, Ayokunle O. Olanrewaju

## Abstract

**Background:** Cytomegalovirus (CMV) can cause severe disease and death in infants and immunocompromised people. Although effective at treating CMV, the first-line drug – ganciclovir (GCV) – has high rates of pharmacokinetic variability which leads to significant rates of underexposure or toxicity. Intracellular concentrations of ganciclovir triphosphate (GCV-TP) – GCV’s active anabolite – are associated with dose-dependent neutropenia and could enable dose individualization to improve treatment efficacy and reduce adverse effects. However, GCV-TP is currently measured using liquid chromatography tandem mass spectrometry (LC-MS/MS) which is impractical for routine use, especially in resource-limited settings, because of its high cost, labor-intensiveness, and need for specialized equipment. To address this gap, we adapted the REverse transcriptase ACTivity with CRISPR (REACTR) assay to measure GCV-TP.

**Methods:** We leveraged our earlier work using REACTR to measure reverse transcriptase (RT) inhibitors used to treat and prevent human immunodeficiency virus (HIV) because GCV-TP serendipitously also inhibits HIV RT. We designed custom DNA templates, primers, and CRISPR complexes to accurately measure GCV-TP spiked into buffer and blood. We evaluated the assay’s analytical performance with 40 dried blood spots from infants with congenital CMV.

**Results:** REACTR reproducibly measured clinically relevant GCV-TP concentrations using a simple workflow and equipment that are readily available in many clinical laboratories. REACTR measurements of clinical samples correlated with LC-MS/MS GCV-TP measurements (r = −0.7891; p<0.0001).

**Conclusions:** This study highlights the potential of REACTR as a rapid and accessible alternative to LC-MS/MS for therapeutic drug monitoring of GCV.

---

Human cytomegalovirus (CMV) is a herpesvirus with an estimated global seroprevalence of over 80% [1]. While infection is typically asymptomatic in immunocompetent individuals, CMV can cause life-long morbidities and even death in immunocompromised or immuno-immature populations like transplant recipients or infants. Congenital CMV (cCMV) affects approximately 1 in 200 newborns, with 10-20% of infants with cCMV exhibiting symptoms at birth [2], [3]. 50% of children with cCMV symptoms at birth suffer from long-term hearing loss, developmental delay, or neurocognitive impairment [4]. Moreover, as many as 10% of infants with symptomatic cCMV die in early infancy due to the disease [3], [5]. Like other herpesviruses, CMV causes an incurable, life-long infection that can become dormant and reactivate over time [6]. While there has been interest in developing CMV vaccines since the 1970s, no CMV vaccine has been licensed to date [7], [8].

Ganciclovir (GCV) and its oral pro-drug valganciclovir (VGCV) serve as the first-line antiviral agents for symptomatic CMV treatment [3]. GCV, or 9-(1,3-dihydroxy-2-propoxymethyl)guanine, is an acyclic guanosine analog that potently inhibits the replication of herpesviruses [9]. Upon cellular uptake, GCV is phosphorylated into its active form, ganciclovir triphosphate (GCV-TP), by viral and host kinases [10]. GCV-TP works by selectively inhibiting extension of viral DNA by herpesvirus DNA polymerases, such as UL54 polymerase [11]. Although effective, GCV therapy is plagued by substantial interindividual variability in drug exposure and high rates of toxicity. With standard dosing, underexposure and inadequate treatment occur in an estimated 30% of infants [12], while overexposure and significant toxicity occur in 25% of infants [13]. These challenges underscore the need for therapeutic drug monitoring (TDM) of GCV to guide personalized dosing.

Current TDM of GCV measures plasma or serum GCV concentrations and is performed using liquid chromatography tandem mass spectrometry (LC-MS/MS) or high-performance liquid chromatography (HPLC) [14]. These tests provide sensitive and accurate GCV measurement but require expensive, centralized equipment and highly trained personnel. Alternative methods for measuring plasma GCV concentrations include Raman spectroscopy, capillary electrophoresis, and electrochemical sensing, but LC-MS/MS remains the gold standard in clinical studies and routine care [14], [15], [16], [17]. Furthermore, clinical studies using LC-MS/MS or HPLC methods for plasma GCV monitoring have presented conflicting evidence for the effectiveness of plasma GCV TDM as a biomarker for treatment optimization [12], [14], [18], [19], [20], [21], [22], [23]. While some studies have reported associations between plasma GCV exposure and treatment outcomes and even target therapeutic ranges for monitoring [19], [20], [24], others have found no such relationship [21], [22], with results varying by study and patient population. As a result, experts have called for further exploration of GCV-TP – GCV’s active intracellular anabolite – as a biomarker for TDM, as it may better predict toxicity and treatment response [22], [25], [26], [27].

A 2015 study demonstrated that intracellular GCV-TP concentrations in kidney transplant recipients were significantly correlated with neutrophil count [22]. In the same study, no associations were seen between plasma GCV or other metabolites of the drug and neutrophil count, suggesting that GCV-TP may be unique in its ability to predict neutropenia. This is the only study of its kind correlating intracellular GCV derivatives with treatment outcomes, likely due to the current cost and complexity associated with GCV-TP measurement. Intracellular GCV-TP concentrations have so far only been measured using LC-MS/MS, a method that requires a complex, multistep procedure to separate GCV-TP from other GCV derivatives before mass spectrometry measurements [28]. This intensive procedure and the increased complexity, time, and equipment associated with GCV-TP measurement make it highly centralized and limit widespread use.

We recently developed the <u>RE</u>verse transcriptase <u>ACT</u>ivity with CRISP<u>R</u> (REACTR) assay for rapid and minimally instrumented measurement of reverse transcriptase (RT) inhibitors used in human immunodeficiency virus (HIV) treatment and prevention [29], [30], [31], [32], [33]. REACTR measures intracellular concentrations of nucleotide analog drugs based on inhibition of DNA synthesis by HIV RT enzyme. Complementary DNA (cDNA) synthesized by HIV RT from a custom DNA template activates a Cas12-crRNA complex, triggering collateral cleavage of DNA reporter molecules. Nucleotide RT inhibitors (NRTIs) compete for incorporation into cDNA with deoxynucleotide triphosphates (dNTPs), such that chain termination, cDNA synthesis, and resulting fluorescence from activated CRISPR complexes are proportional to the NRTI concentration in the reaction. GCV-TP, despite canonically targeting herpesvirus DNA polymerases, also inhibits HIV RT via delayed chain termination [34], [35]. Consequently, while REACTR was originally developed for measuring HIV medications, GCV-TP’s inhibition of HIV RT allowed us to adapt REACTR to measure GCV-TP.

In this study, we adapted the REACTR assay for measuring GCV-TP in buffer and dried blood spots (DBS) and evaluated the assay’s analytical performance with DBS from a clinical pharmacokinetic (PK) trial. We first designed custom DNA templates and optimized assay conditions for detecting GCV-TP spiked into buffer and blood matrices, characterized its sensitivity and reproducibility, and then validated its performance using clinical DBS samples from the Randomized Controlled Trial of Valganciclovir for Cytomegalovirus Infected Hearing Impaired Infants (ValEAR) study (ClinicalTrials.gov #NCT03107871) [36]. Finally, we compared REACTR GCV-TP measurements with those obtained by LC-MS/MS to assess concordance with the gold-standard method.

## RESULTS AND DISCUSSION

### Adapting REACTR for measuring GCV-TP

REACTR uses a ssDNA template with a primer binding region followed by a repeat region specific to the nucleotide analog drug of interest, a complementary crRNA recognition region, a PAM sequence, and a poly T tail (**Figure 1A**). To adapt REACTR to measure GCV-TP, a guanine analog, a cytidine repeat region was incorporated into the template. Assays were run with the DNA template, a reverse primer, RT, dNTPs, a Cas12a-crRNA complex specific to the DNA template, DNA reporters with fluorophore-quencher pairs, and sample containing GCV-TP (**Figure 1B, Table S1**). As in previous demonstrations of REACTR, as the amount of drug in the sample increases, DNA chain termination increases [33]. This prevents synthesis of the crRNA recognition region and results in reduced fluorescence (**Figure 1B**). Thus, we can infer GCV-TP drug concentrations based on REACTR assay fluorescence.

**Figure 1.**
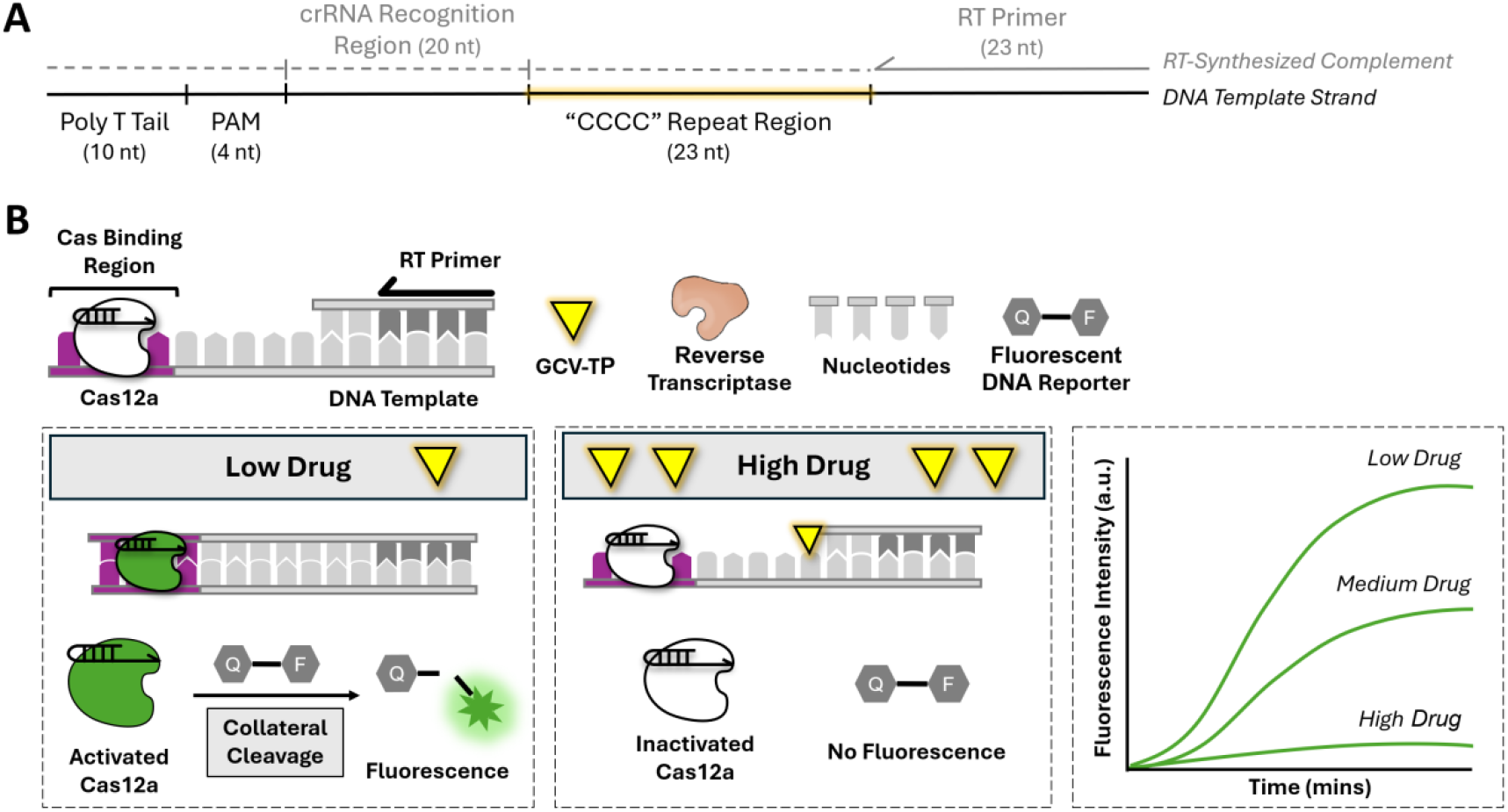
Overview of the REverse transcriptase ACTivity with CRISPR (REACTR) assay for measuring GCV-TP based on its inhibition of DNA synthesis by HIV RT enzyme. (**A**) The REACTR template optimized for measuring GCV-TP (a guanine analog) includes a primer binding region followed by a cytidine repeat region, a complementary crRNA recognition region, a PAM sequence, and a poly T tail. (**B**) At low drug concentrations, HIV RT can synthesize full-length DNA with CRISPR binding regions that activate Cas12a complexes that cleave DNA reporters leading to fluorescence. Conversely, at high drug concentrations, DNA chain termination occurs and inactivated Cas12a complexes are unable to generate fluorescence.

### REACTR can measure clinically relevant GCV-TP concentrations in buffer

We assessed REACTR’s ability to measure clinically relevant GCV-TP concentrations by spiking the drug into aqueous buffer at concentrations from 0.1 mM to 0.01 nM and added the spiked solutions into REACTR assays. Real-time measurements showed decreased fluorescence as GCV-TP concentration increased (**Figure 2A, Figure S1A**). We explored the role of template design on REACTR’s sensitivity to GCV-TP by running the assay with a template containing a “CCCC” repeat region designed to bind GCV-TP (a guanine analog) and a “TTTA” repeat region originally designed for tenofovir diphosphate (an adenine analog used in HIV treatment) (**Figure 2B, Figure S1B**). The “CCCC” template had a 50% Inhibitory Concentration (IC_50_) (95% Confidence Interval [CI]) of 10.8 nM (9.1 to 12.9 nM), while the “TTTA” template required at least 3 orders of magnitude greater GCV-TP concentrations before inhibition was observed and did not provide a quantifiable IC_50_ within the span of concentrations tested. These results, in agreement with our past work, show that REACTR assay parameters like DNA template sequence can be tuned to specifically measure clinically relevant levels of different antiviral drugs.

**Figure 2.**
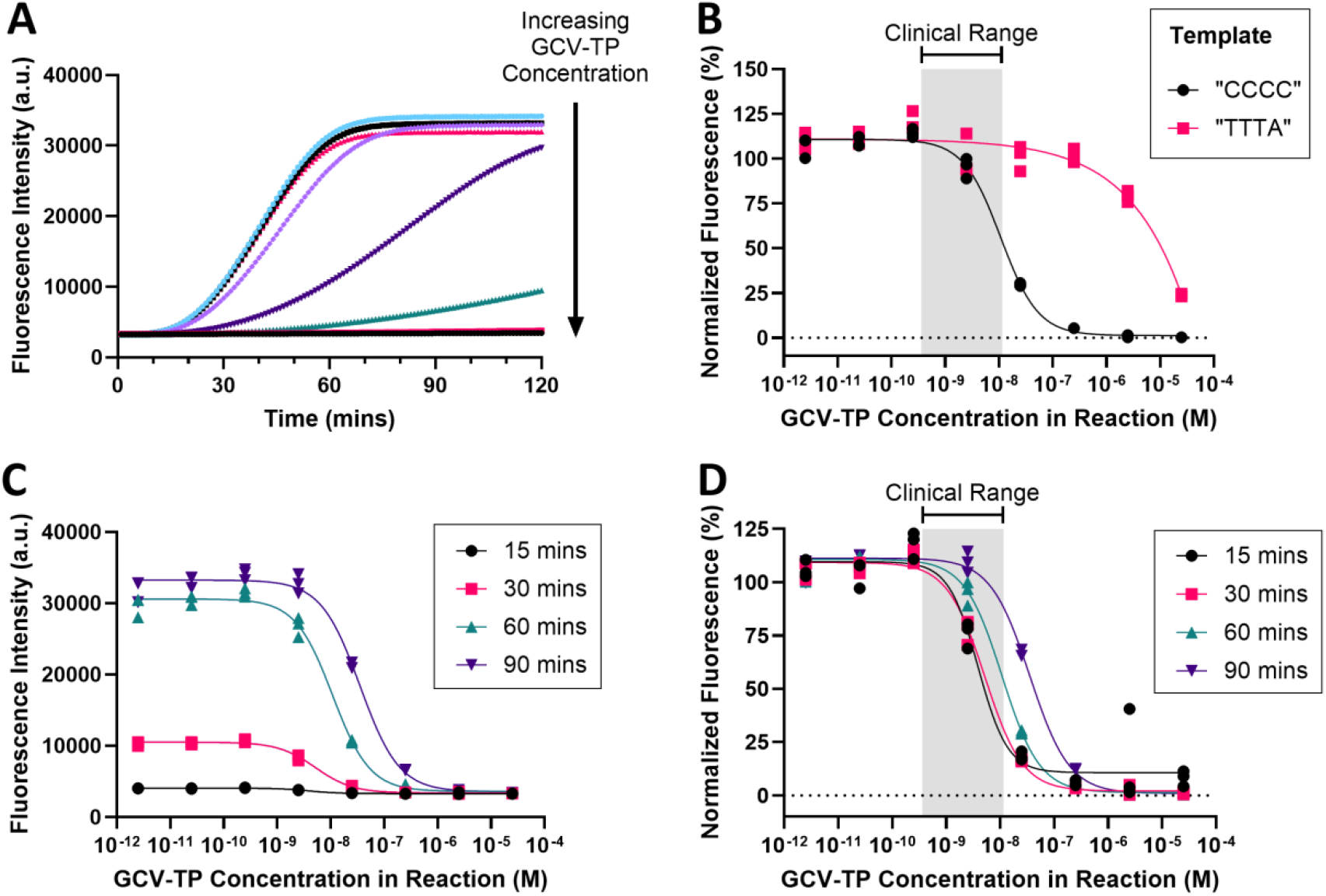
REACTR sensitively and specifically measures varying concentrations of GCV-TP spiked in buffer. (**A**) Mean real-time REACTR curves using “CCCC” templates with varying concentrations of GCV-TP spiked in buffer. (**B**) Normalized RT inhibition curves for GCV-TP in REACTR assays using templates with either “CCCC” or “TTTA” repeat regions show specificity in detecting the guanine analog drug using the cytidine-enriched template. (**C**) RT inhibition curves for GCV-TP at various timepoints – extracted from raw data in (A) – show an increase in fluorescence signal as assay incubation time increases. (**D**) Normalized RT inhibition curves for GCV-TP at various timepoints show that shorter assay incubation times enable detection of lower GCV-TP concentrations. n = 3 technical replicates. Gray regions indicate the clinical range of GCV-TP (0.362 to 11.5 nM) in DBS of infants with cCMV.

We also explored the role of reaction incubation time on REACTR assays with GCV-TP by generating enzyme inhibition curves at the 15-, 30-, 60-, and 90-minute time points (**Figure 2C**). As expected, raw fluorescence signal increased as assay incubation time increased. Normalizing data showed that the IC_50_ (95% CI) of GCV-TP at 15, 30, 60, and 90 min were 3.9 nM (N/A to 6.5 nM), 5.2 nM (4.3 to 6.4 nM), 10.8 nM (9.1 to 12.9 nM), and 35.4 nM (29.9 to 42.6 nM), respectively (**Figure 2D**). Thus, there is a tradeoff between the assay fluorescence signal intensity and the ability to detect low concentrations of GCV-TP.

### REACTR accurately measures clinically relevant GCV-TP concentrations spiked in DBS

We next adapted REACTR to measure GCV-TP in DBS as a convenient way to test archived clinical samples. Using DBS from healthy donors not receiving any antiviral drugs, we developed a simple sample preparation workflow that used thermolabile proteinase K (TPK) and heat to denature blood proteins (**Figure 3**). GCV-TP was spiked into blood eluted from DBS at concentrations from 0.1 mM to 0.01 nM prior to the addition of TPK and heat. The dilution and protein denaturation steps in the sample preparation process reduce non-specific inhibition of HIV-RT by blood matrix components including hemoglobin and immunoglobulins.

We found that, even with our sample preparation process, the maximum signal intensity (from “no drug” controls) and the assay variation increased with REACTR in DBS compared to buffer (**Figure S2**). Although in Figure 2D a 30-minute incubation time with REACTR in buffer provided inhibition curves that enabled GCV-TP quantification within the clinical range, we chose a 60-minute assay time with REACTR in DBS to improve assay performance. At a 30-minute incubation time, the decrease in the maximum signal intensity with REACTR in DBS compared to REACTR in buffer was 46.2% and the coefficient of variation (CV) of “no drug” controls in DBS was 8.8% compared with 4.4% in buffer (**Figure S2B-D**). Meanwhile, at a 60-minute incubation time, the maximum signal intensity decreased by 40.7% in DBS compared to buffer, and the CV of “no drug” controls was 3.7% in DBS compared with 3.1% in buffer (**Figure 4A**).We attribute these differences in performance between REACTR in DBS and buffer to incomplete elimination of blood matrix effects with our relatively simple sample preparation process. Nevertheless, we prioritized assay simplicity and the use of instruments that are typically available in most clinical laboratories (vortexer, heat block, microcentrifuge, and fluorescence reader) for all our assay preparation and readout. Despite the decrease in raw signal intensity, the normalized inhibition curve for GCV-TP spiked in DBS closely mirrored that seen in buffer at 60 minutes, with a modest rightward shift in IC_50_ between buffer (10.8 nM, 95% CI = 9.09 to 12.9 nM) and blood (21.2 nM, 95% CI = 15.6 to 28.8 nM) (**Figure 4B**).

**Figure 3.**
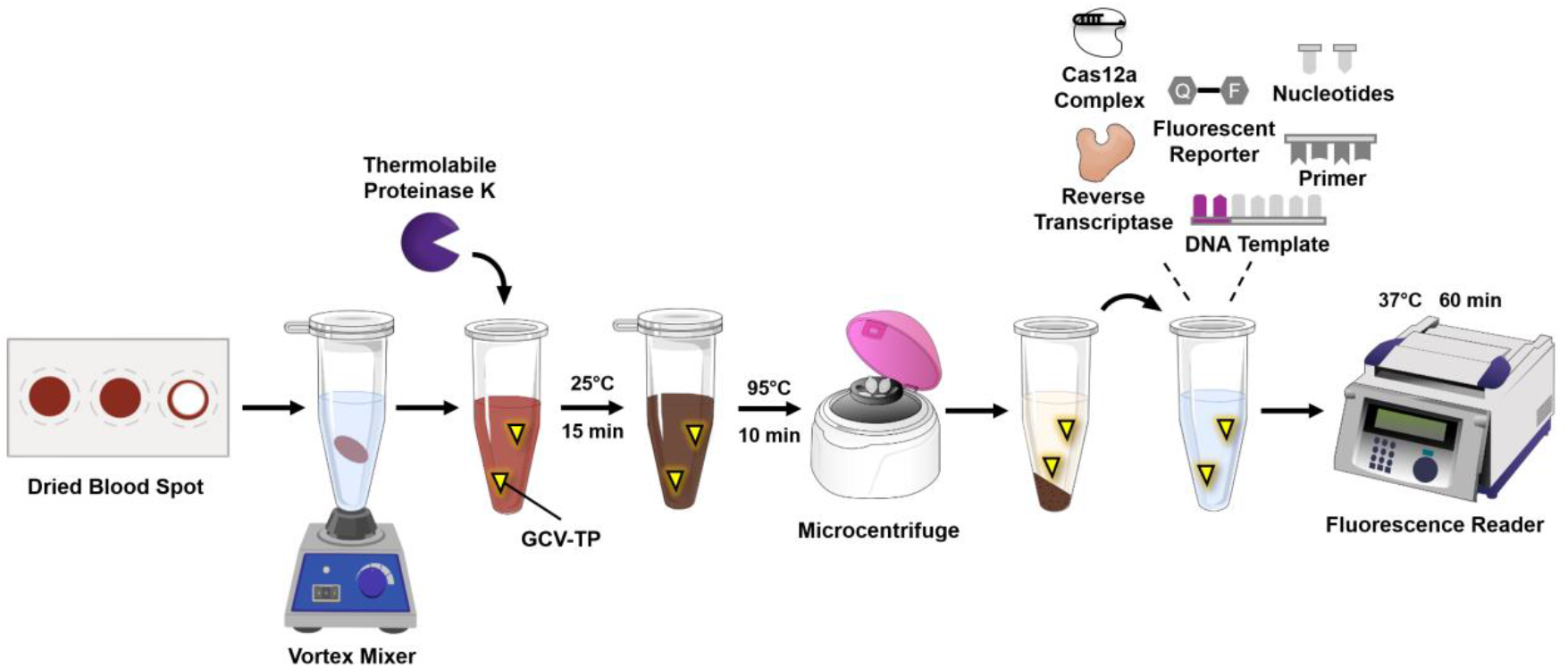
Simple sample preparation workflow for measuring antiviral drugs in DBS using the REACTR enzymatic assay. DBS punches are vortex mixed in water to lyse red blood cells and release intracellular nucleotide analogs, then treated with thermolabile proteinase K to digest blood proteins, followed by heat denaturation. The supernatant with extracted GCV-TP is then added to a tube with RT, DNA template, primer, Cas12a complex, fluorescent reporter, and nucleotides, then incubated at 37°C for 60 minutes in a fluorescence reader.

**Figure 4.**
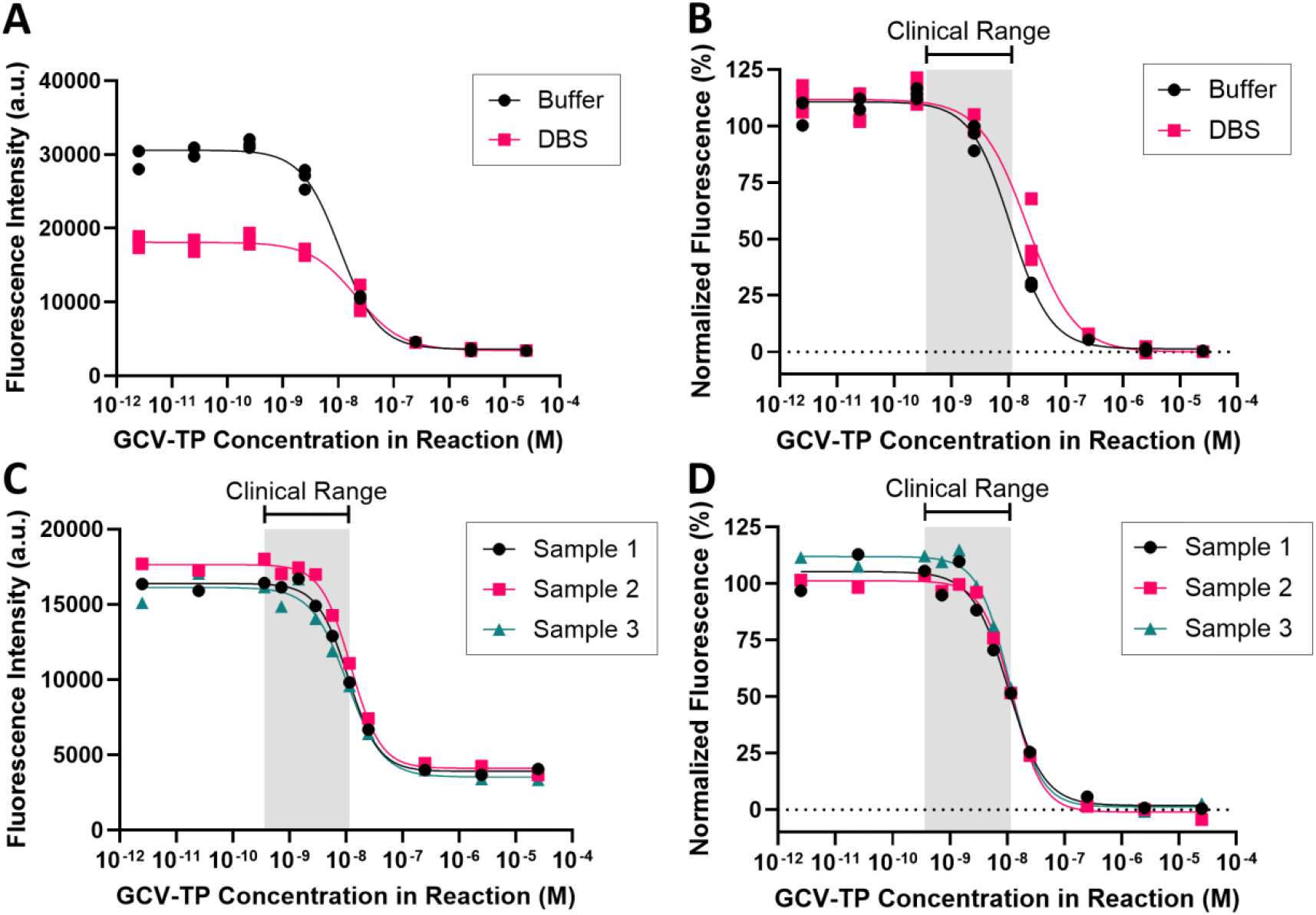
REACTR provides consistent results with GCV-TP spiked in DBS. (**A**) RT-inhibition curve with varied GCV-TP concentrations shows significantly lower maximum fluorescence signal with spiked DBS compared with spiked buffer at a 60-minute assay incubation time. n = 3 technical replicates. (**B**) Normalized inhibition curve for GCV-TP spiked in buffer and DBS – extracted from data in (A) – showing excellent agreement between buffer and blood with a slight rightward shift for the assay in DBS. (**C**) RT inhibition curves for GCV-TP spiked in blood from 3 independent blood samples show excellent agreement across samples. Each point on the curve represents the mean of 3 technical replicates. (**D**) Normalized 60-minute RT inhibition curves for GCV-TP spiked in 3 independent blood samples – extracted from data in (C) – show excellent consistency across samples.

We evaluated sample-to-sample variation using spiked DBS samples generated from independent blood samples not containing any antiviral drugs (**Figure 4C-D, Figure S3**). Both raw fluorescence curves (**Figure 4C**) and normalized curves (**Figure 4D**) were highly consistent across samples. Across the three samples, the average intra-assay CV was 12.8% and the inter-assay CV was 10.6%. We also calculated the Z’ score to assess the assay’s ability to distinguish between positive and negative controls and as an indicator of its suitability for high throughput screening [42]. The average intra-assay Z’ score was 0.54 and the inter-assay Z’ score was 0.56 – both above the quality cutoff of 0.5 for Z’ value typically used in high-throughput screening assays.

We also evaluated the assay’s ability to distinguish drug concentrations in the clinical range from the “no drug” control. Of the sampled concentrations within the clinical range, the raw fluorescence became significantly different from “no drug” control at the 5.8 nM drug condition (p = 0.0214) when assessed with a one-way ANOVA with multiple comparisons. The same cutoff was seen after normalizing fluorescence (p = 0.0001). This indicates that the assay in its current form can detect GCV-TP concentrations at the upper end of the clinical range (0.362 to 11.5 nM GCV-TP) in DBS, which may be useful for detecting potentially toxic concentrations of GCV-TP in blood.

### REACTR measurements correlate with LC-MS/MS-measured GCV-TP concentrations

To assess performance of the REACTR assay with clinical samples, we tested 40 DBS samples collected from 16 infants enrolled in the ValEAR trial with GCV-TP concentrations ranging from 0 to 1891 fmol/6 mm punch. **Figure 5** shows REACTR fluorescence intensity measurements compared with LC-MS/MS GCV-TP concentrations from the ValEAR study. REACTR measurements of the clinical DBS samples correlated with LC-MS/MS measurements of GCV-TP (n = 40, Pearson r = −0.7891; p < 0.0001; 95% confidence interval (CI) = −0.8834 to −0.6332), with a coefficient of determination (r^2^) of 0.6226. LC-MS/MS correlations to raw fluorescence values and normalized values (**Figure S3**) were highly consistent. These initial results demonstrate the potential of REACTR to provide accurate GCV-TP measurements at a fraction of the time, cost, and complexity required for LC-MS/MS testing.

**Figure 5.**
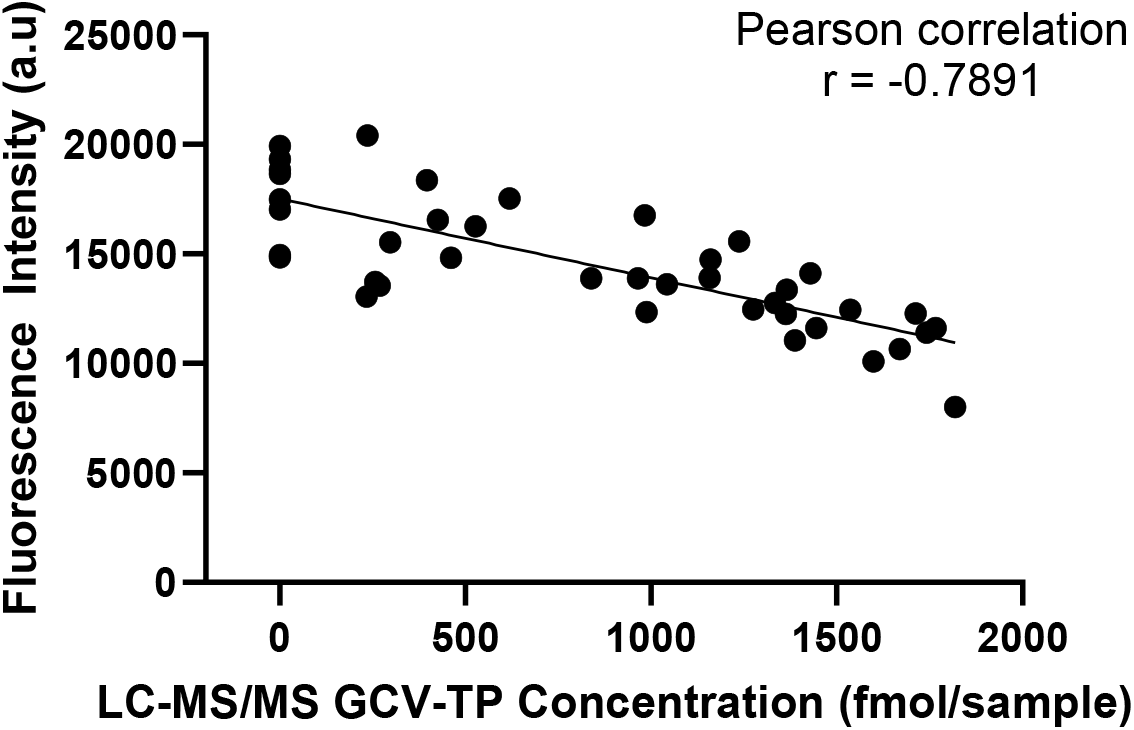
REACTR results correlate with LC-MS/MS GCV-TP concentrations in DBS from a clinical pharmacokinetic trial. n = 40 DBS from infants with cCMV. Each point represents the mean of 3 technical replicates.

### Limitations and future directions

While this study demonstrates the feasibility of adapting REACTR for intracellular GCV-TP monitoring, several limitations exist. First, the current assay is limited to detecting GCV-TP levels at the upper end of the clinical range. Improving assay sensitivity through reagent optimization and enhanced sample preparation or readout methods will be important to expand detection to the lower end of the clinical range. We previously developed and validated a mathematical model showing that increasing DNA template length, increasing the number of potential chain termination events by enriching for the complement of the drug in the DNA template, and decreasing the concentration of competing nucleotides are all effective strategies for increasing the analytical sensitivity of assays like REACTR [31]. Switching from using HIV RT to the UL54 polymerase, the canonical target of GCV-TP, may also improve the assay’s sensitivity to lower concentrations of the drug. Exploring alternate sample preparation or readout strategies that reduce the influence of blood matrix effects without sacrificing assay simplicity would also help to improve the performance of the assay. Second, clinical validation was performed using DBS from a relatively small cohort of infants with cCMV. Although REACTR measurements correlated with LC-MS/MS, larger studies will be necessary to establish assay reproducibility and to correlate GCV-TP levels with treatment response and toxicity outcomes. Third, the optimization and clinical validation shown here focused exclusively on infants with cCMV. GCV is also used widely in transplant recipients who may exhibit different pharmacokinetic profiles and thus may have a different clinical range. In future work we hope to validate our assay for measuring GCV-TP in DBS from transplant populations to broaden clinical utility.

## Conclusion

This study validates the REACTR platform for measuring intracellular GCV-TP, the active metabolite of GCV used in the treatment of CMV infection. By leveraging GCV-TP’s ability to inhibit HIV RT and coupling changes in cDNA synthesis by HIV RT to Cas12a-based detection, REACTR measures clinically relevant concentrations of GCV-TP in DBS samples in 60 minutes. REACTR was highly reproducible in DBS across independent blood samples, with average intra-assay and inter-assay CVs of 12.8% and 10.6%, respectively, and intra-assay and inter-assay Z’ scores above 0.5. Importantly, REACTR measurements of clinical DBS samples from infants living with cCMV correlated with LC-MS/MS measurements of GCV-TP (r = −0.7891; p < 0.0001), demonstrating agreement with the current gold-standard analytical method. Compared to LC-MS/MS, REACTR offers several advantages: (1) streamlined sample preparation, (2) reduced turn-around time, and (3) use of commonly available and relatively inexpensive laboratory equipment. These features could enable expanded access to therapeutic drug monitoring of GCV-TP, which has been called for but not widely implemented due to the limitations and associated costs/complexities of current techniques. Future work will prioritize improving assay sensitivity through reaction chemistry and workflow optimization and conducting larger clinical validation studies across both cCMV and transplant populations. Taken together, our results demonstrate the feasibility of REACTR as a rapid, accessible, and cost-effective approach for monitoring GCV-TP levels with the potential for use in clinical trials and/or routine care.

## MATERIALS AND METHODS

### REACTR assay workflow

REACTR workflows were adapted from a previously described method by Singh et al. [33]. The Cas12a-crRNA complex was created by incubating 100 nM LbCas12a (M0653T, New England Biolabs) and 100 nM crRNA (Integrated DNA Technologies) in REACTR buffer containing 50 mM Tris-HCl (77-86-1, Sigma-Aldrich) at pH 8, 50 mM Potassium Chloride (KCl) (60142-100 ML-F, Sigma-Aldrich), 15 mM Magnesium Chloride (MgCl2) (7786-30-3, Sigma-Aldrich), 7 mM dithiothreitol (DTT) (3483-12-3, Sigma-Aldrich), and 0.06% Triton X-100 (9036-19-5, Sigma-Aldrich) for 15 minutes at 25°C.

For spiked buffer experiments, GCV-TP (NC2174868, Axxora) was diluted in nuclease-free water at varying concentrations from 0.1 mM to 0.01 nM. 40 µL REACTR assays containing 10 µL of sample or drug dilution, 25 nM Cas12-crRNA complex, 25 nM dNTPs, 0.5 nM DNA template, 12.5 nM primer, 500 nM fluorescent reporter (Integrated DNA Technologies), and 0.04 units/µL of HIV RT (LS005009, Worthington Biochemical) in REACTR buffer were added to 96-well armadillo PCR plates (Thermo Scientific, AB2396). dNTP and template concentrations were chosen to maximize sensitivity to GCV-TP via decreased competition of dNTPs while maintaining signal intensity [31]. All assays were run against “no-drug” positive controls and “no-enzyme” negative controls. Reactions were incubated at 37°C in a thermal cycler (Bio Rad CFX96 Touch Real-Time PCR Detection System) for 60 to 120 minutes. Fluorescence measurements were taken every minute during the incubation period. Raw and analyzed data is available on Zenodo [37].

### DBS sample preparation

6 mm DBS punches were collected in 1.7 mL Eppendorf tubes and eluted with 67.5 µL of nuclease free water. The tube was then vortexed for 1 minute and centrifuged in a microcentrifuge for 30 seconds. Eluted blood was transferred to a separate tube and thermolabile proteinase K (P8111L, NEB) was added to create a final concentration of 25% blood and 2.5% proteinase K. The tube was gently mixed before being incubated at 25°C for 15 minutes followed by a heat inactivation step at 95°C for 10 minutes. The tube was centrifuged again in a microcentrifuge for 1 minute to pellet proteinaceous material. The supernatant was then added to the REACTR assay.

For spiked DBS experiments, GCV-TP was diluted in nuclease-free water (H_2_O) at varying concentrations from 1 mM to 0.1 nM and added to eluted DBS punches at a ten-fold dilution prior to the proteinase K. The elution volume in spiked DBS experiments was reduced accordingly to maintain a final concentration of 25% blood and 2.5% proteinase K. Remaining sample preparation was performed as described above. Sample to sample variation was assessed using three independent blood samples. Sample 1 was derived from pooled blood purchased from BioIVT, and samples 2 and 3 were from independent donors not receiving GCV.

### Clinical DBS sample collection

Authentic clinical DBS samples were obtained from the ValEAR trial, a randomized, multicenter, placebo-controlled clinical trial of VGCV in infants with a positive CMV test [38]. The trial was conducted under central institutional review board (IRB) approval (University of Utah/Primary Children’s Hospital board; cIRB_00090760, NCT03107871) and in accordance with the Declaration of Helsinki. Infants were enrolled with informed parental consent and randomized to receive VGCV (16 mg/kg twice daily for 6 months) or placebo. Monthly venipuncture blood samples were collected for PK analyses. DBS samples were stored at –80°C until analysis.

### LC-MS/MS Analysis of GCV-TP

LC-MS/MS quantification of GCV-TP in DBS samples is described by Lindquist-Kleissler et al. and adapts previously described methods [36], [39], [40], [41]. Briefly, each 6 mm DBS punch was extracted into 0.5 mL of 70% methanol (MeOH) in H_2_O by sonication for 15 minutes. GCV-TP in the supernatant was fractionated from other phosphorylated species using Waters QMA solid phase extraction (SPE) cartridges, followed by dephosphorylation with acid phosphatase and cleanup using a Phenomenex Strata-X SPE cartridge. A 30 μL aliquot of each sample was then injected onto a TSQ Vantage triple quadrupole mass spectrometer (ThermoScientific) interfaced with an Accela UHPLC pump and autosampler (ThermoScientific). Chromatographic separation was achieved under isocratic flow of 0.1% formic acid in 99% H_2_O and 1% MeOH at 0.250 mL/min through a 100 x 2 mm Polar RP analytical column (Phenomenex^®^) equipped with a 4 x 3 mm Security Guard Polar-RP™ column (Phenomenex^®^). Tandem mass spectrometry was operated in positive electrospray ionization (ESI) mode, monitoring mass transitions of 256.1 to 152.1 for GCV and 261.2 to 152.1 for the internal standard. Quantification was based on a calibration curve fit with a 1/x^2^ weighted linear regression over the range of 50 to 4000 fmol/sample. Intra-day accuracy was within ±4.0% of nominal values and the coefficient of variation (CV) was within 4.9% across three runs with 6 technical replicates each of quality control samples.

### Clinical range estimate

The clinical range used to guide assay development was derived from the range of concentrations observed in the ValEAR study (233 to 1891 fmol/6 mm punch). Due to the low sample size of participants on valganciclovir used in this dataset, we extended this range to 100 to 3200 fmol/6 mm punch, based on the assumption that variability at the upper end of the range would exceed that of the lower end [22]. Assuming that each 6 mm punch holds 17.2 µL of sample based on the storage capacity of Whatman™ 903 Proteinsaver cards, we converted values in fmol/6mm punch to molar concentrations in REACTR. The resulting molar clinical range within the REACTR assay was 0.362 nM to 11.6 nM, accounting for a final blood dilution of 6.25% in the reaction.

### Data analysis and statistical methods

Enzyme inhibition curves were generated in GraphPad Prism 10 using REACTR fluorescence values at the 60-minute timepoint. When applicable, fluorescence intensities were normalized to the average of the “no drug” positive and “no RT” negative controls as maximum and minimum values, respectively. Curves were fit to four-parameter logistic regression curves. The intra-assay coefficient of variation (CV) was calculated from the baseline-subtracted mean fluorescence of the positive control from 3 technical replicates. The inter-assay CV was calculated from the baseline-subtracted mean fluorescence of the positive control from 3 independent replicates, each representing the mean of 3 technical replicates. Intra-assay Z-prime (Z’) score was calculated using the raw fluorescence mean and standard deviation values of assay controls from 3 technical replicates. Inter-assay Z-prime (Z’) score was calculated using the raw fluorescence mean and standard deviation values of assay controls from 3 independent replicates, each representing the mean of 3 technical replicates. All clinical samples were run and analyzed blinded. One-way ANOVA tests with multiple comparisons were performed using GraphPad Prism 10 and the reported p-values were adjusted to account for multiple hypothesis tests. Correlation between LC-MS/MS GCV-TP concentrations and fluorescence were determined using a Pearson’s correlation coefficient.

## Acknowledgments

***Acknowledgements***. We would like to credit DBCLS for the centrifuge and thermal cycler icons in Figure 2, which are licensed under CC-BY 4.0 Unported (https://creativecommons.org/licenses/by/4.0/) and Servier for the microtube icons in Figure 2, which are licensed under CC-BY 3.0 (https://creativecommons.org/licenses/by/3.0/) and had the contents and lids modified.

## Author Contributions

**W.A.C**.: Conceptualization; Formal analysis; Investigation; Methodology; Validation; Writing – original draft; Writing – review & editing; Visualization; Project administration. **M.A.S**.: Conceptualization; Methodology; Writing – review & editing. **C.H.L**.: Methodology. **M.M.C**.: Funding acquisition; Methodology; Writing – review & editing. **C.A.C**.: Methodology. **A.H.P**.: Resources. **J.E.R**.: Resources; Writing – review & editing. **A.O.O:** Conceptualization; Funding acquisition; Methodology; Project administration; Supervision; Resources; Writing – review & editing.

## Data Availability

Nucleotide sequences of REACTR reagents are available in supplementary Table 1. Raw and analyzed data are available on Zenodo [37].

## Financial Support

This work was supported by the Arnold and Mabel Beckman Foundation and the Washington Entrepreneurial Research Evaluation and Commercialization Hub (WE-REACH) to A.O.O.

### Abbreviations

GCV: Ganciclovir
VGCV: Valganiclovir
GCV-TP: Ganciclovir triphosphate
TDM: Therapeutic drug monitoring
REACTR: REverse transcriptase ACTivity with CRISPR
LC-MS/MS: Liquid chromatography tandem mass spectrometry
HPLC: high-performance liquid chromatography
RT: Reverse transcriptase
cCMV: Congenital cytomegalovirus
NRTI: Nucleotide reverse transcriptase inhibitor
DBS: Dried blood spots

