## Supplementary Information for "REverse-transcriptase ACTivity with CRISPR (REACTR) Assay for Rapid and User-Friendly Therapeutic Drug Monitoring in Cytomegalovirus Care"

### SUPPLEMENTARY MATERIAL

**Table 1. Custom DNA and RNA sequences used in REACTR assay.**

| Name | Sequence (5’ to 3’) |
|---|---|
| 80nt “CCCC” DNA template | TTTTTTTTTTTTTGATGATGTGAAGGTGTTGTCGCCCCCCCCCC CCCCCCCCCCCCCCTATCTTTCCTCTTAATTCGACG |
| 80nt “TTTA” DNA template | TTTTTTTTTTTTTGATGATGTGAAGGTGTTGTCGTTTATTTATTT ATTTATTTATTTCTATCTTTCCTCTTAATTCGACG |
| DNA reverse primer | CGTCGAATTAAGAGGAAAGATAG |
| crRNA target region | TACTACACTTCCACAACAGC |
| crRNA | rUrArArUrUrUrCrUrArCrUrArArGrUrGrUrArGrArUrArUrGrArUrGrUr GrArArGrGrUrGrUrUrGrUrCrG |
| Fluorescent reporter | /56-FAM/TTATT/3IABkFQ/ |

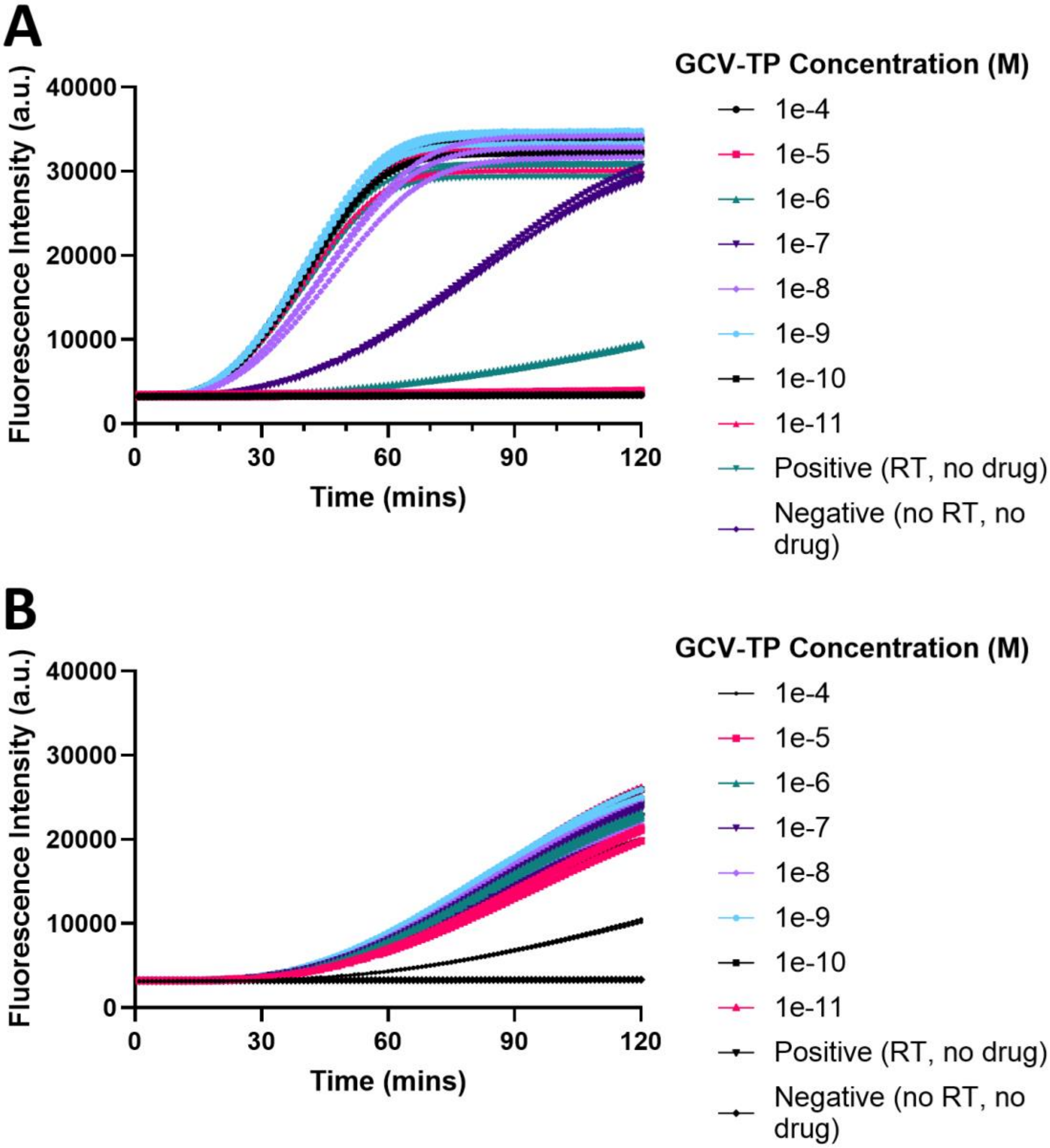


**Figure S1. Real time REACTR curves in buffer.** (**A**) Real-time REACTR curves using "CCCC" templates with varying concentrations of GCV-TP spiked in buffer show enzyme inhibition starting around 1e-7 M, indicating pronounced inhibition of DNA synthesis by the guanine analog drug due to Watson-Crick-Franklin base pairing. (**B**) Real-time REACTR curves using "TTTA" templates with varying concentrations of GCV-TP spiked in buffer only show enzyme inhibition around 1e-4 M, indicating significantly delayed inhibition with the thymidine-rich template. These results, in agreement with our past work, demonstrate the specificity of tuning the REACTR assay for different nucleotide analog drugs. n = 3 technical replicates.

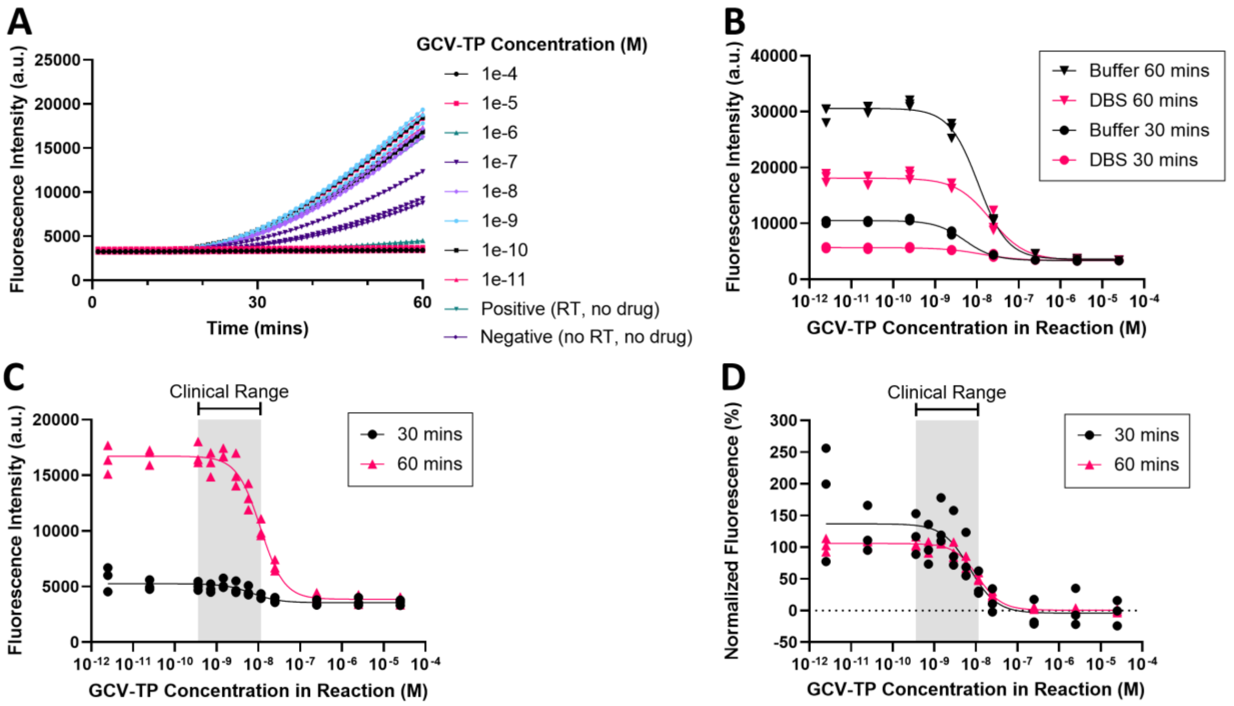


**Figure S2. REACTR optimization in spiked DBS samples shows differences between buffer and DBS and improved performance in DBS with 60 min assay time.** (**A**) Real-time REACTR curves with varying concentrations of GCV-TP spiked in DBS sample 1. **(B)** RT-inhibition curves with varied GCV-TP concentrations spiked in buffer or blood at 30-minute and 60-minute assay incubation times. n = 3 technical replicates. (**C**) RT-inhibition curves for GCV-TP-spiked DBS at 30-minute or 60-minute assay incubation times from 3 independent DBS samples. Each point on the curve represents the mean of 3 technical replicates. (**D**) Normalized RT-inhibition curves for GCV-TP-spiked DBS at 30-minute or 60-minute assay incubation times from 3 independent DBS samples. Each point on the curve represents the mean of 3 technical replicates.

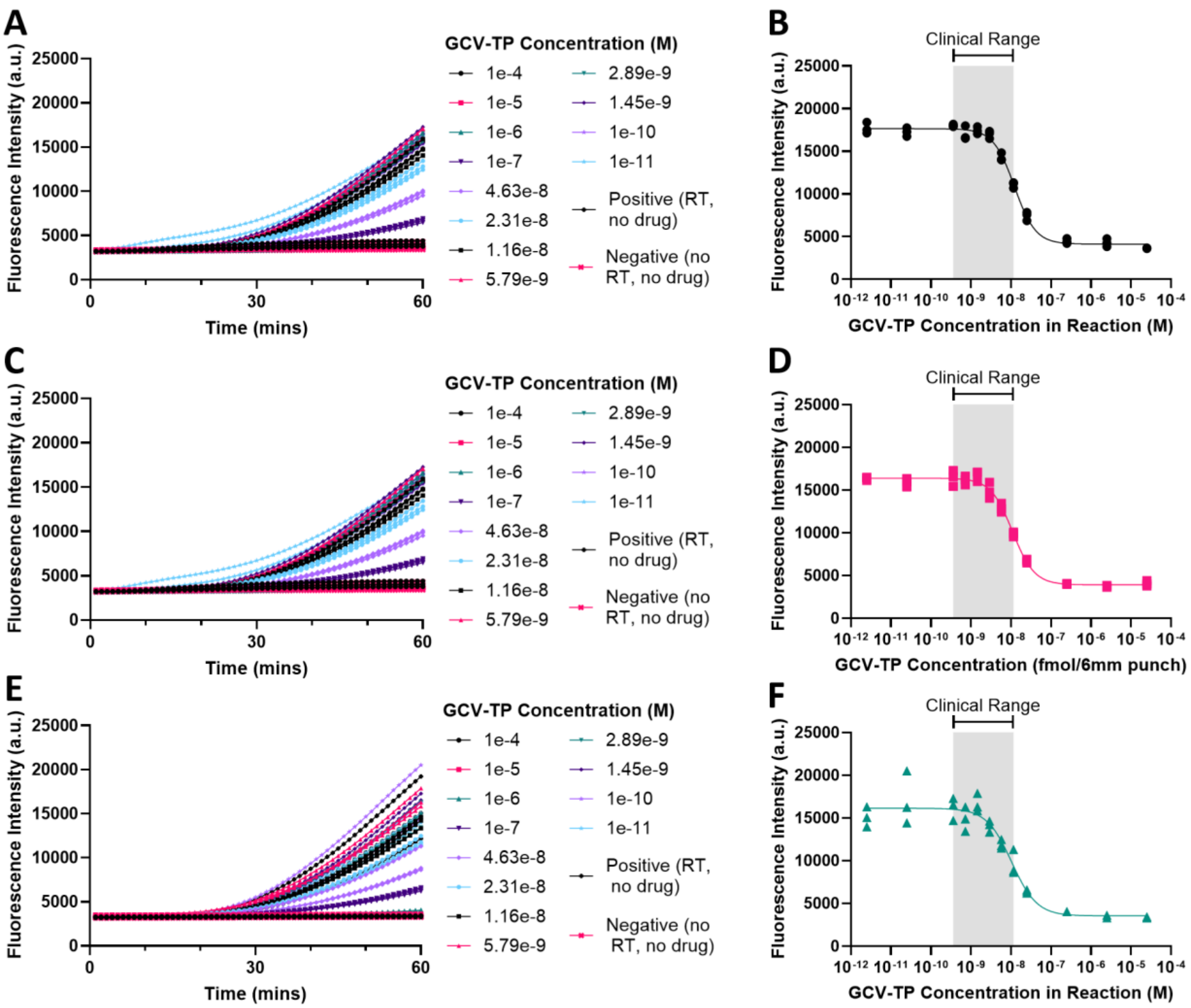


**Figure S3. REACTR with GCV-TP spiked in 3 independent DBS samples shows consistency across all three samples. (A)** Real-time REACTR curves with varying concentrations of GCV-TP spiked in DBS sample 1. (**B**) 60-minute RT inhibition curve for GCV-TP-spiked DBS from sample 1. (**C**) Real-time REACTR curves with varying concentrations of GCV-TP spiked in DBS sample 2. (**D**) 60-minute RT inhibition curve for GCV-TP-spiked DBS from sample 2. (**E**) Real-time REACTR curves with varying concentrations of GCV-TP spiked in DBS sample 3. (**F**) 60-minute RT inhibition curve for GCV-TP-spiked DBS from sample 3. n = 3 technical replicates.

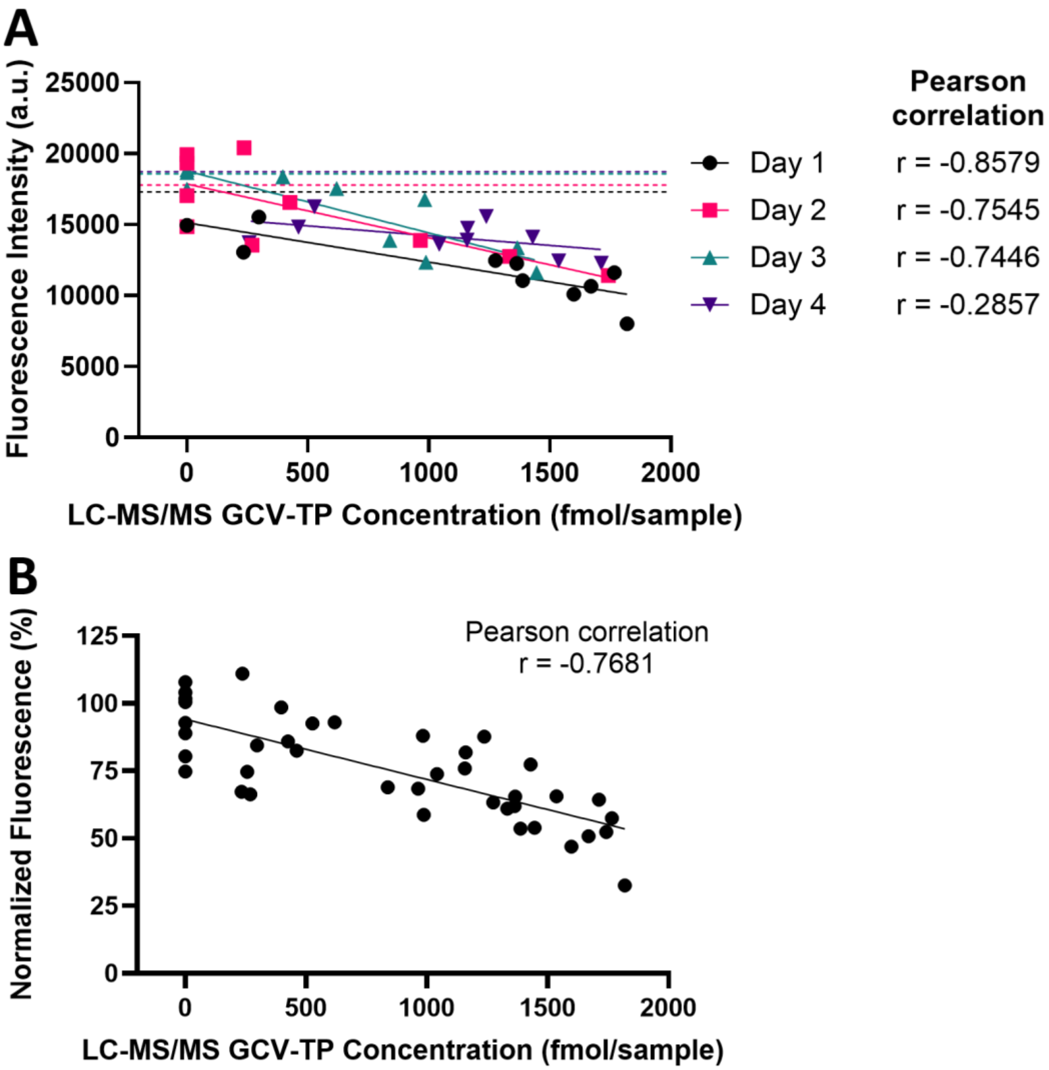


**Figure S4. REACTR correlation with LC-MS/MS GCV-TP concentrations in DBS from a clinical pharmacokinetic trial.** (**A**) Correlation of REACTR fluorescence and LC-MS/MS GCV-TP measurements by day. 10 samples were run per day in technical triplicate. The dotted line represents the mean fluorescence of the BioIVT pooled blood no drug control. (**B**) Correlation of REACTR Normalized fluorescence and LC-MS/MS GCV-TP measurements. n = 40 DBS from infants with cCMV. Each data point represents the mean of 3 technical replicates.
